# Ultrasound-activated drug release with extracellular vesicles

**DOI:** 10.1101/2025.08.11.669710

**Authors:** Xi Shi, Weilong He, Haiyue Xu, Leonardo Clark, Helena M. Zeng, Ashwin Gupta, Nitya Mirle, Lief Fenno, Huiliang Wang

## Abstract

Precise, noninvasive targeting and controlled drug release to the nervous system could transform both basic research and clinical therapy. In this work, we developed a focused ultrasound–responsive delivery system based on extracellular vesicles (EVs), which are naturally occurring and biocompatible phospholipid nanocarriers. To enable the ultrasound responsiveness, a lipid–gas phase transition material Perfluoropentane (PFP) was co-encapsulated with therapeutic agents into EVs via low-temperature sonication. The resulting EVs had a size of 100–200 nm and demonstrated efficient, on-demand release upon ultrasound stimulation. In primary cultured neurons, EVs loaded with lidocaine successfully suppressed calcium activity and showed minimum cytotoxicity. Finally, we demonstrated *in vivo* application of this system for modulation pain sensitivity in rats. Our EVs-drugs platform demonstrated a safe, and ultrasound-activated drug delivery with high spatiotemporal control, showing strong potential therapeutic applications.

## 1. Introduction

Focused ultrasound, when coupled with artificial drug carriers, has demonstrated noninvasive drug release capabilities with high spatial and temporal precision.^1^ Ultrasound-responsive drug carriers include microbubbles, lipid nanoparticles, nanoemulsions, polymers and hydrogen-bonded organic frameworks (HOFs), which can either be inherently responsive to ultrasound or incorporating of sonosensitizers.^1–5^ For example, ultrasound-triggered drug release has been achieved using carriers and sonosensitizers such as IR780 and protoporphyrin IX.^6,7^ The sonosensitizers would generate active oxygen after applying ultrasound which would destroy the lipid membrane resulting drug release. In addition, certain lipid nanoparticles can release drugs through thermal activation due to their low lipid melting points.^8^ Upon ultrasound generating slight heat would dissolve the membrane and release drug. We have recently developed HOFs to achieve ultrasound-programmable drug release for sono-chemogenetics.^3^ However, all these chemically synthesized carriers might suffer from limitations such as cytotoxicity.^9,10^

In this work, we leveraged extracellular vesicles (EVs) to develop an ultrasound-triggered drug release system. EVs are naturally secreted by many cell types, with size ranging from 40 nm to 100 nm, and have been investigated as drug carriers for central nervous system applications since the 2010s.^12^ Perfluorocarbons, including perfluoropentane (PFP), have been FDA-approved and have demonstrated safety and efficacy in preclinical studies for targeted drug delivery and disease diagnosis, particularly in cancer and cardiovascular diseases.^11,13^ Inspired by previous work using ultrasound-responsive, perfluorocarbon-based lipid nanoemulsions for ultrasound-triggered drug release,^2^ we have employed sonication to co-load EVs with perfluorocarbons and therapeutic agents, successfully enabling the ultrasound-triggered drug release with our PFP-encapsulated EV systems. As a demonstration for application of this system for pain management, we encapsulated local anesthetic drug lidocaine in our system and used it for both *in vitro* inhibition of cultured neuronal activity and *in vivo* modulation of pain threshold in the sciatic nerve in rats.

## 2. Results

### 2.1. Characterization of isolated EVs and EVs loaded with drugs

To develop a safer ultrasound-activated drug release system, we utilized EVs derived from HEK293T cells as carrier vehicles and perfluoropentane (PFP) as a liquid-to-gas phase-change material. (**Figure 1a**). EVs were isolated from the conditioned media (2.5% exosomes (EVs)-depleted FBS) of HEK293T cells using differential centrifugation followed by ultracentrifugation.^14^ Cryo-electron microscopy (Cryo-EM) revealed that the EVs exhibited a spherical morphology with well-defined membranes. (**Figure 1b**, left). Furthermore, cryo-tomography of the EVs showed the presence of cellular proteins both on the membrane surface and within the vesicles (**Figure S1**). In order to determine the size of EVs, nanoparticle tracking analysis (NTA) were used to measure the size of isolated EVs to be approximately 100 nm (**Figure 1c**). To encapsulate both drugs and PFP, we explored methods commonly used for hydrophilic molecule encapsulation in EVs, including electroporation (as used in cells) or sonication (as used in liposomes).^15^ To evaluate encapsulation efficiency, Rhodamine B, a hydrophilic fluorescent dye, was used as a model compound. While encapsulation in either electroporation or sonication method worked in a small 1 mL system,^16^ sonication was ultimately chosen for its scalability of encapsulation in a 10 mL encapsulation system. The resulting PFP and dye loaded EVs (EVs-drugs) exhibited approximately a 1.5-to-2-fold increase in size compared to native EVs, as determined by both Cryo-EM image (**Figure 1b**, right) and NTA analysis (**Figure 1c**). Notably, EVs-drugs showed a spherical shape with a denser core, suggesting successful encapsulation of PFP and Rhodamine B dye (**Figure 1b**, right).

**Figure 1.**
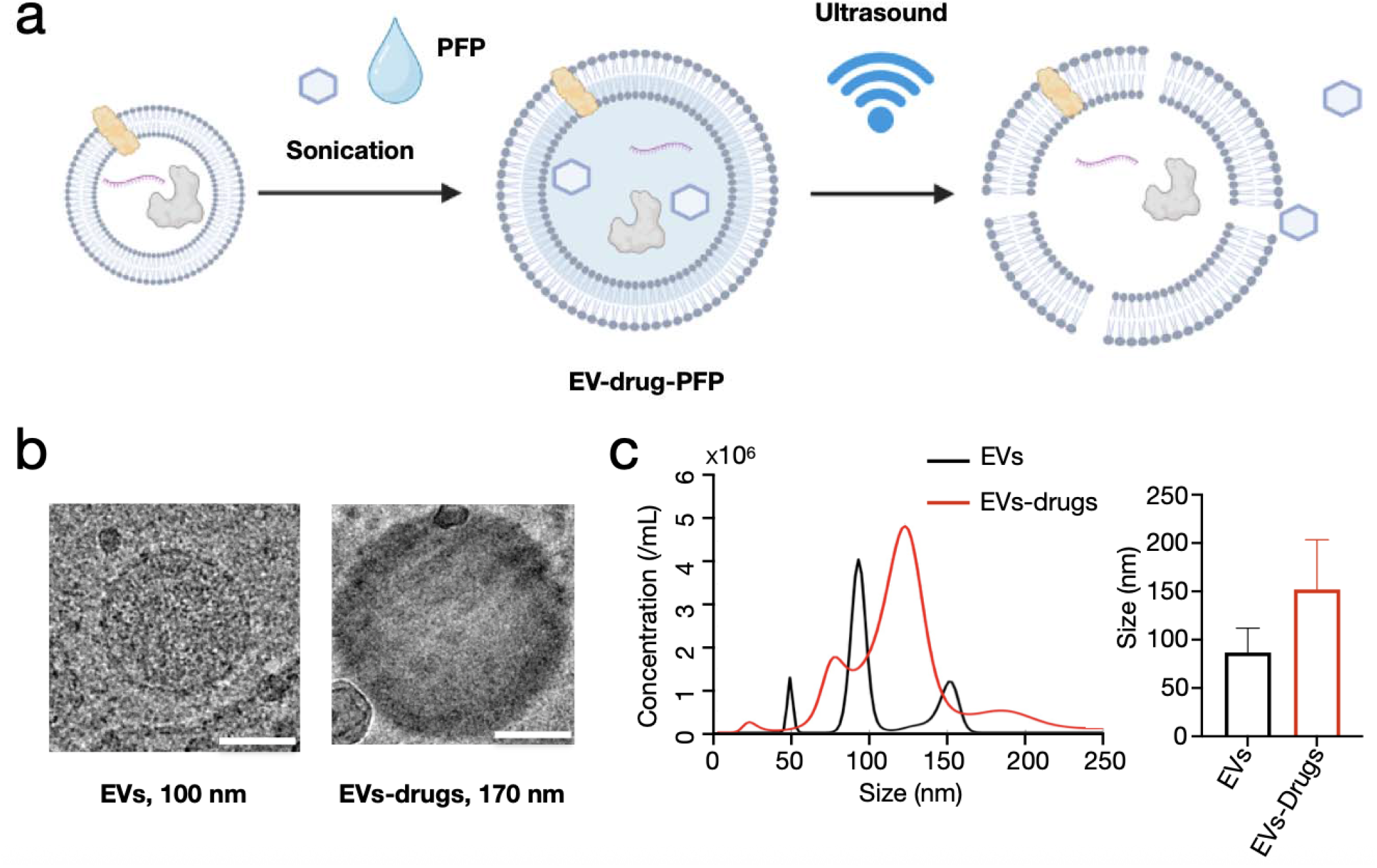
Schematic illustration and characteristics of EVs isolation and drug loading. (**a)** The schematic illustration of EVs loading drugs and ultrasound-activated drug release. (**b**) Cryo-EM images of EVs and EVs-drugs and their sizes. Scale bar, 50 nm. (**c**)The concentration and size distribution of EVs and EVs-drugs measured by nanoparticle tracking analysis.

### 2.2. *In vitro* ultrasound-activated drug release

To determine the drug release from our EVs-drug formulation using ultrasound, we employed rhodamine B dye and PFP encapsulated EVs. After sonicating the EVs with rhodamine B and PFP, we performed three to five rounds of low-speed centrifugation (2,000 × g) using pre-cooled PBS to wash the encapsulated particles. Red-colored pellets were observed, indicating the successful loading of rhodamine B into the EVs. The loading efficiency of rhodamine B was calculated to be approximately 4–5%, based on a standard calibration curve of the free dye.

To evaluate ultrasound-triggered drug release, we applied focused ultrasound (1.5 MHz transducer) with a 25% duty cycle, and a 5 Hz burst frequency for 5 minutes, using varying amplitude settings. The amount of dye released was quantified by measuring the absorbance of rhodamine B in the supernatant using UV-vis spectroscopy (**Figure 2a**). To determine the total encapsulated dye, EVs-drugs were heated to 65□°C for 5–30 minutes to ensure complete release, after which no pellet remained post-centrifugation. The absorbance spectrum of the resulting supernatant was measured by UV-Vis again to quantify total dye loading.

**Figure 2.**
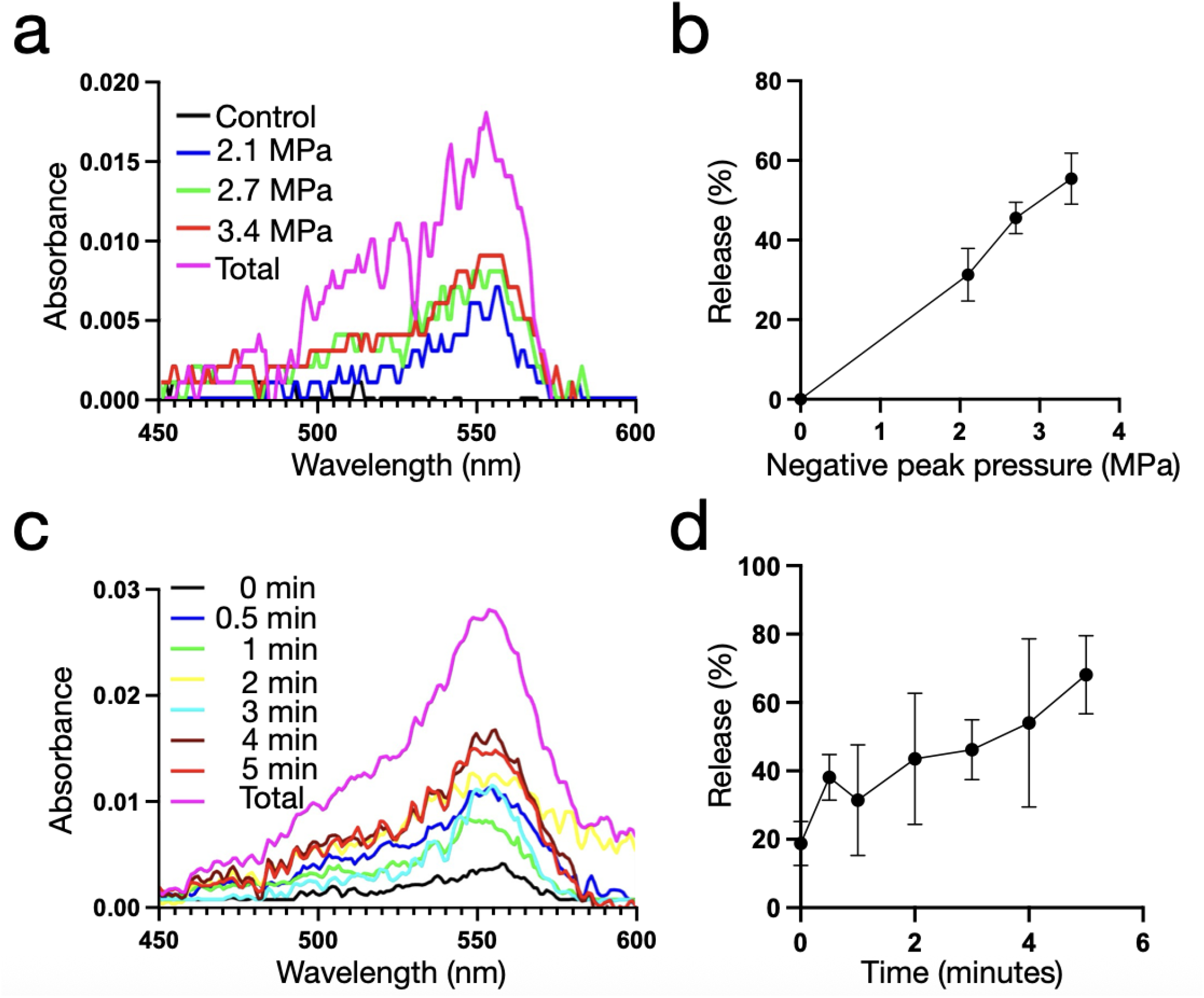
*In vitro* ultrasound-activated drug release with EVs-drugs. (**a**) Absorption Spectrum of Rhodamine B (RhB) release with different ultrasound negative peak pressures (1.5 MHz ultrasound for 5 minutes with 25% duty cycle and 5 Hz). The control sample is the supernatant of the washed EVs-drugs. Total release sample is EVs-drugs heated under 65 °C. (**b**) Drug release efficiency under different ultrasound negative peak pressure. (n=3, mean ± SEM) (**c**) Absorption Spectrum of Rhodamine B (RhB) release with different ultrasound application times (1.5 MHz transducer and 2.1 MPa peak pressure), (d) Accumulation drug release from 0 to 5 minutes every minute. (n=3, mean ± SEM)

Release efficiency increased with higher ultrasound negative peak pressures (**Figure 2b**). For example, a 100% amplitude setting—corresponding to a negative peak pressure of 3.4 MPa—resulted in approximately 60% dye release. In addition to ultrasound pressure, the duration of exposure also influenced release efficacy. At a 2.1 MPa peak pressure, increasing the sonication time from 30 seconds to 5 minutes led to progressively higher dye release efficiency (**Figure 2c&2d**). These results demonstrate that our EVs-drug system can be activated by ultrasound for controlled drug release, with both ultrasound pressure and exposure time modulating the amount released.

### 2.3. *In vitro* neuromodulation with ultrasound-activated drug release from EVs

We tested neuronal inhibition using EVs encapsulated with local anesthetic drug lidocaine. Lidocaine acts as a sodium channel blocker, suppressing ion influx and thereby inhibiting neuronal activity.^17^ (**Figure 3a**) To evaluate ultrasound-activated drug release for modulating neuronal activity, cultured neurons were transduced with AAV9-hSyn-GCaMP6s-WPRE-SV40 to express GCaMP6s— a genetically encoded calcium indicator that will change its green fluorescence with calcium influx, thereby indicating the changing neuronal firing (**Figure 3b**). EVs-drug in PBS (40 μL) were added to one well containing GCaMP6s-expressing neurons, which was then filled with a pre-warmed neuron culture medium and placed on the microscope. A convex water-coupled ultrasound transducer (without gas in the bubble), covered by an ultrasound-protective membrane, was positioned in contact with the medium in the imaged well.

**Figure 3.**
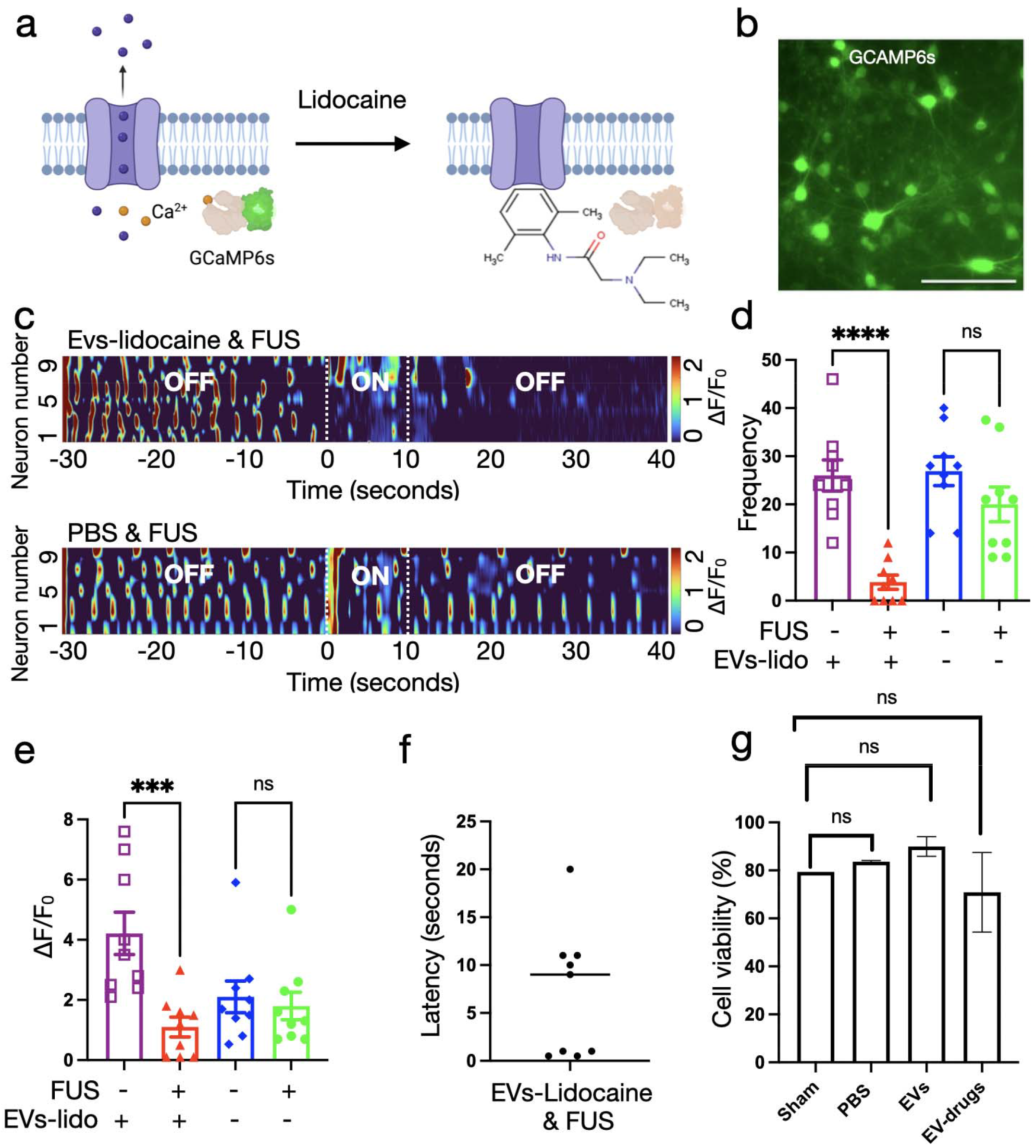
*In vitro* neural modulation with ultrasound-activated EVs drug release. (**a**) The illustration of lidocaine binding to the ion channel. (**b**) Primary cultured neurons expressed GCaMP6s after being transduced with AAV9-hSyn-GCaMP6s-WPRE-SV40 for four to seven days. Scale, 50 μm. (**c**) Heatmap of neurons inhibition with the application of 10 seconds of ultrasound (1.55 MPa) triggered EVs-lidocaine drug release (n=9, mean ± SEM). (**d**) Neuronal firing frequency before and after focused ultrasound (FUS), P<0.0001. (**e**) Changing fluorescence of neurons before and after FUS with or without EVs-lidocaine drug release. P<0.001. (**f**) The latency of neuron inhibition after ultrasound EVs-lidocaine drug release. (**g**) The neuron toxicity of EVs-drugs after 24h incubation showed no difference with control groups (Sham or PBS). (n=3, mean ± SEM)

Before ultrasound stimulation, neurons incubated with EV-drugs exhibited spontaneous neuronal activity, indicating that the encapsulated drugs remained inactive. Upon 10 seconds of focused ultrasound (1.55 MPa, 1.5 MHz) application, we observed rapid inhibition of neuronal activity. Neurons treated with EV-lidocaine and FUS displayed a marked reduction in calcium signaling (**Figure 3c**). The neuronal firing frequency decreased after EVs-lidocaine and FUS activating lidocaine to block spontaneous activity of neurons (**Figure 3d**), while the control neurons with FUS but not EVs-lidocaine showed no difference in activity frequency. There was also a significant decrease in changing fluorescence (dF/F0) after FUS induced the drug release from EVs-lidocaine, which suggested neuron inhibition (**Figure 3e**). Approximately 70% of these neurons ceased spontaneous activity within 10 seconds of FUS stimulation (**Figure 3f**), with suppression lasting for 40 to 80 seconds. These results demonstrate that our EV-drug-PFP platform enables rapid and efficient drug release upon FUS stimulation and can effectively modulate neuronal activity for durations exceeding one minute.

Next, we evaluated the cytotoxicity of EV-drug formulations in cultured cortical neurons. Cell viability was assessed using propidium iodide (PI), which stains the nuclei of dead cells red, and Hoechst 33342, a blue, fluorescent dye that labels all cell nuclei, allowing for total cell count. We used EVs encapsulating both PFP and lidocaine in our cytotoxicity assays. Each well of a 48-well plate containing cultured neurons received 0.8 × 10□ EV-drugs or control EVs in 40□μL of PBS. After incubation, neurons were stained with PI and Hoechst 33342, washed three times with Neurobasal medium, and immediately imaged using epi-fluorescence microscopy. The results indicated no significant difference in neuronal viability between EV-drug-treated and control groups at 24h (**Figure 3g** and **Figure S2**). These findings suggest that EV-drugs have minimum non-toxicity to neurons.

### 2.4. *In vivo* ultrasound-activated drug release from EVs to block sciatic nerve

Next, we evaluated drug release from EVs-lidocaine triggered by focused ultrasound (FUS) in a rat sciatic nerve model. This model is relevant for assessing analgesia, as numbness of the sciatic nerve results in decreased pain sensitivity in the hind paw. EVs-lidocaine was first injected into the muscle tissue adjacent to the sciatic nerve, followed by a 5-minute diffusion period to allow the EVs-lidocaine to localize near the nerve (**Figure 4a**). Subsequently, FUS was applied to the shaved skin overlying the sciatic nerve for 8 minutes (1.33 MPa, 1,5MHz and 50% duty in 1Hz pulse). Pain sensitivity in the hind paw was assessed at 30 minutes after injecion, applied between the fourth and fifth toes, and the responses were compared to baseline mechanical thresholds (**Figure 4b**).

**Figure 4.**
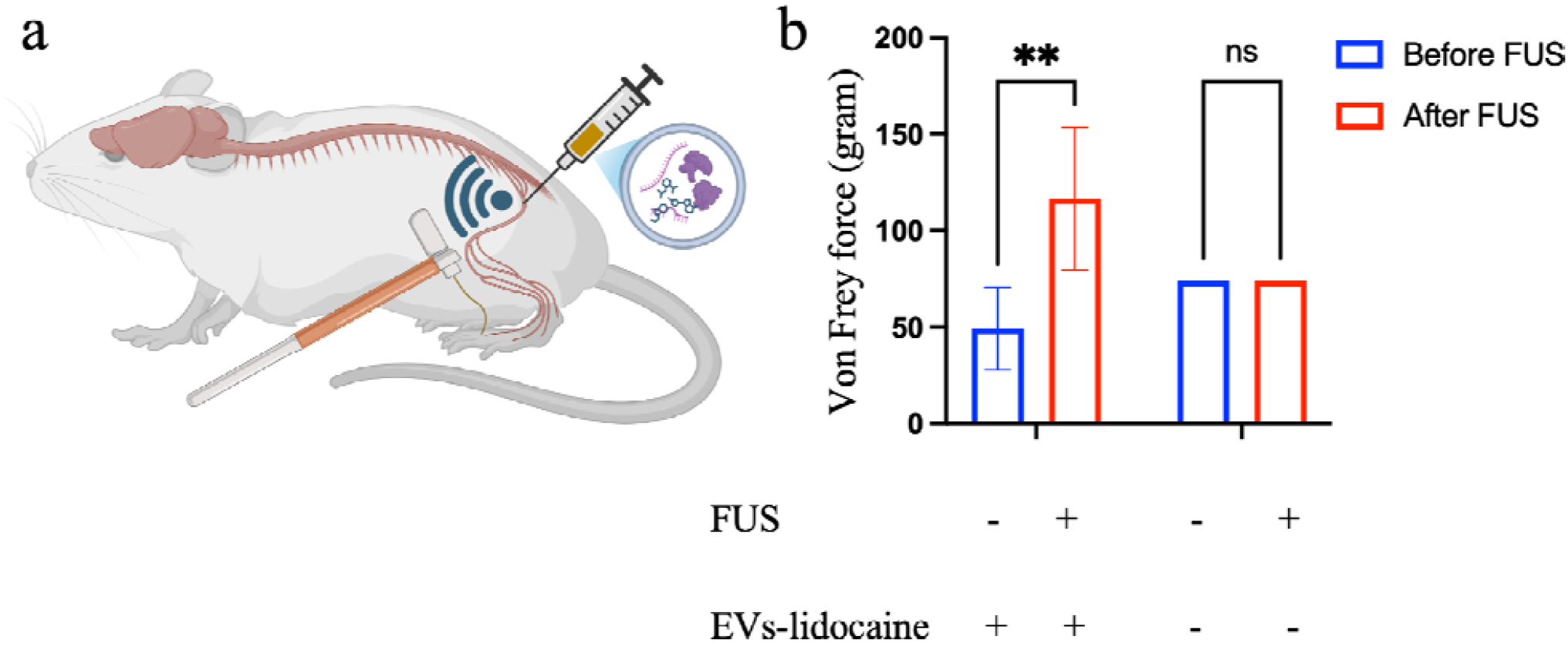
*In vivo* EV-drugs release in rats’ sciatic nerves. (**a**) Illustration of EV-drugs injecting to the sciatic nerve and FUS activation of drug release with the von Frey tests at the toe of the rats. (**b**) von Frey force showed significant nerve block after EV-lidocaine injection and FUS activation compared with the control rats with PBS injection and FUS. (n=3, mean ± SEM)

The results showed that application of FUS to EVs-lidocaine increased paw withdrawal thresholds significantly from ∼40 grams to ∼140 grams (over 100% increase in pain threshold). These findings suggest that the FUS activated EVs-lidocaine can effectively increase pain sensitivity in sciatic nerve. However, no change in pain threshold in control rats with only the application of FUS but without EVs.

## 3. Summary

We have demonstrated that extracellular vesicles (EVs) encapsulated with perfluoropentane (PFP) serve as an effective FUS-triggered drug release platform. By co-encapsulating PFP and hydrophilic molecules, we achieved effective drug release upon FUS stimulation. Also, we characterized drug release efficiency across varying ultrasound pressures and found both increased ultrasound pressure and increased ultrasound application length could increase the amounts of release drugs. Next, we demonstrated ultrasound-activated EVs-drugs release for inhibition of neuronal activity with EVs-Lidocaine in primary cultured neurons. To assess biocompatibility, we exposed neurons to high concentrations of EVs-Lidocaine and observed no significant cytotoxicity with EVs-drugs. Finally, we applied ultrasound-activated EVs-lidocaine *in vivo* and demonstrated effective modulation of pain sensitivity of sciatic nerve in rats. In summary, the FUS-triggered EVs-drugs platform represents a promising and safe strategy for on-demand drug delivery to the neurons, with strong translational potential for clinical applications.

## 4. Experimental methods

### Cell culture and EVs isolation

Human embryonic kidney (HEK) 293T cells were cultured in high-glucose Dulbecco’s Modified Eagle Medium (DMEM) supplemented with 10% fetal bovine serum (FBS), 1× penicillin-streptomycin (P/S), and 2 mM L-glutamine in 182 cm^2^ flasks (25 mL media per flask) at 37□°C in a 5% CO□ incubator until reaching confluency. Cells were passed at a 1:3 ratio and cultured in media containing 2.5% exosome-depleted FBS, P/S, and L-glutamine for 60–72 hours. Typically, 45 to 50 flasks were used to yield ∼10^12^ EVs. Cell culture supernatants were collected and centrifuged at 500 × g for 10 min and then at 2,000 × g for 10 min to remove cells and debris. The supernatant was filtered using a 0.22 µm PES membrane and concentrated by tangential flow filtration (TFF) with a 100 kDa molecular weight cutoff. The final 100– 200 mL of concentrated supernatant was ultracentrifuged at 100,000 × g for 1 hour at 4□°C to isolate EVs.

### Cryo-EM imaging

EVs at a concentration of 10^12^ particles/mL in PBS were used for cryo-electron microscopy (cryo-EM) imaging. Samples were applied to Lacey Carbon Film-coated 200 mesh Cu grids with a continuous ultrathin carbon layer (SKU: LC200-Cu-CC-25, EMS). A lacey carbon film grid (Electron Microscopy Sciences, Hatfield, PA) was plasma-cleaned for 1 minute in a PelcoeasiGlow glow discharge system (Ted Pella Inc., Redding, CA). 4 uL of each exosome sample (i.e., different drug loaded accordingly) was deposited on the grid and excess volume were blotted away for 3.5 seconds using grade 595 vitrobot filter paper (Ted Pella Inc., Redding, CA) with a force of -4 at 25 °C in a 100% humidity chamber. The grids were then plunged frozen into liquid ethane using a Vitrobot Mark IV (Thermo Scientific, Waltham, Massachusetts). Frozen grids were imaged on a Thermo Fisher Scientific Glacios (Thermo Scientific, Waltham, Massachusetts) operated at 200 kV and equipped with a Falcon 4 direct detector (Thermo Fisher Scientific). Imaging was conducted at the Sauer Structural Biology Laboratory, a core facility at The University of Texas at Austin. Image processing was performed using CryoSPARC software.

### Nanoparticle tracking analysis (NTA)

NTA was performed using the NanoSight NS500 and ViewSizer 3000 (HORIBA) instruments at the Allen J. Bard Center for Electrochemistry, University of Texas at Austin. EV samples were serially diluted to a final concentration of 1–3 × 10□ particles/mL. The mode particle size was used for plotting data.

### Drug encapsulation in EVs

To encapsulate drugs, 0.3 mL of perfluoropentane (PFP) was mixed with a saturated drug solution (e.g., lidocaine, 20 mg/mL Rhodamine B, or L-glutamate) in 5 mL PBS. Five milliliters of isolated EVs were added and gently mixed by pipetting. The mixture was sonicated in an ice-cold water bath for 30 seconds, with tube mixing every 10 seconds, followed by a 2-minute incubation on ice. This cycle was repeated six times. The resulting EVs–drugs nanoparticles were washed three times by centrifugation at 2,000 × g for 10 minutes at 4□°C. The final EVs–drugs pellet was either stored at –80C°□ or used immediately.

### Ultrasound EVs-drugs release

Drug release from EV–rhodamine B was tested using a 1.5 MHz concave transducer. EVs-rhodamine B samples (100–500 µL) were freshly mixed and loaded onto the cleaned transducer surface. Focused ultrasound (FUS) was applied (1–5 minutes, 25% duty cycle, 5 Hz). Post-sonication, samples were transferred to PCR tubes and centrifuged at 2,000 × g for 10 minutes at 4□°C. Supernatants (2.5 µL) were collected carefully without disturbing pellets and analyzed via UV-vis spectrophotometry by measuring absorbance between 200–800 nm, focusing on the Rhodamine B peak at ∼560 nm. Total Rhodamine B content was determined by heating EVs-Rhodamine B samples at 65□°C for 5-30 minutes or several hours until no pellet remained after centrifugation.

### Cell viability assay

Primary cortical neurons were isolated from embryonic day 14 mouse cortices and cultured in Neurobasal medium supplemented with L-glutamine, P/S, and B-27 supplement. On day two, glial growth was inhibited using glial inhibitors. Neurons were used for viability assays on day four. 40 μL of EVs (0.8 × 10^9^ particles), EV-lidocaine (0.8 × 10^9^ particles), or PBS were added per well in 48-well plates. After 24 hours without media change, neuron viability was assessed using 5 µg/mL propidium iodide (PI) and Hoechst 33342 (1:2,000 dilution of a 10 mg/mL stock) staining at 37⍰°C for 5 minutes. Neurons were then washed three times with pre-warmed Neurobasal medium, replaced with fresh full medium, and immediately imaged using epifluorescence microscopy. Eight fields per well were captured and analyzed with Fiji/ImageJ to quantify red (dead) and blue (total) cells. Each condition included ≥1,000 neurons. Data were analyzed using Prism software, and statistical significance was determined via two-way ANOVA.

### Calcium imaging of neural activity

Primary cortical neurons were transduced with AAV9-hSyn-GCaMP6s-WPRE-SV40 and incubated for around a week to express the green GCaMP6s. Then 40 μL of EVs-lidocaine (0.8 × 10□ particles) were added to one well of the 48-well plates primary neurons. Fill the well with prewarmed Neurobasal media and imaging with epi-fluorescence microscopy (Leika, DMi8) with 20 X magnification and water bubble transducer mounted in touch with the media of shooting neurons. Focused ultrasound setting is 1.55 MPa, 1.5 MHz for 10 seconds. Images were taken every 100 ms for 70 seconds (700 frames) and plotted the ΔF/F0 in Fiji/ImageJ software. The heatmap of neural inhibition was plotted with MATLAB.

### In vivo neuromodulation in healthy Sprague-Dawley (SD) rats

All animal procedures were designed and taken according to the National Institutes of Health Guide for the Care and Use of laboratory animals, approved by the Institutional Animal Care and Use Committee at the University of Texas at Austin (AUP-2023-00306) and were supervised and supported by the Animal Resources center (ARC). 5-7 weeks of male Sprague-Dawley (SD) rats were purchased from Charles River and were housed in ARC and experimentally taken when rats aged at 8-12 weeks.

EVs-lidocaine ultrasound-activated drug release *in vivo* determination was measured by von Frey assay. SD rats were trained for two days before von Frey assay. On the day of von Frey, we determined the base line at the digits of 4 and 5 in healthy untreated SD rats. Then the SD rats were shaved and injected with 1 mL of EVs-lidocaine or PBS (control groups) under light isoflurane anesthesia (1-2%) to the muscle near the sciatic nerve with 0.8 to 1 cm depth. Waiting for five minutes for the EVs-lidocaine diffused to the sciatic nerve and applied focused ultrasound (FUS) for eight minutes (1.33 MPa, 1.5 Hz, 50 % duty of 1Hz pulse). Von Frey assay was taken 30 minutes after FUS of those injection rats.

## Acknowledgment

We gratefully acknowledge the Center for Electrochemistry, University of Texas at Austin, for providing research support for this work through the CEC Instrumentation Facility maintained under Grant no. H-F-0037 from the Welch Foundation. We thank Dr. Jaso McLelan shared tangential flow equipment and Dr. Cristopher Sullivan shared ultracentrifuge equipment. We also thank Dr. Axel Brilot for cryo-EM data analysis, as well as Dr. Zunlong Ke for taking and analyzing the cryo-tomography. Prof. Huiliang Wang acknowledges funding support from the National Science Foundation (NSF) CAREER award (2340964), NIH Maximizing Investigators’ Research Award (National Institute of General Medical Sciences 1R35GM147408), UT Austin Portugal Exploratory Research Projects Grant, Alpha-1 Foundation Pilot and Feasibility Grant, Robert A. Welch Foundation Grant (No. F-2084-20240404) and Craig H. Neilsen Foundation Pilot Research Grant. We acknowledge BioRender.com for some figure drawing and MATLAB for the data plotting.

## Author contributions

Conceptualization, X.S., W.H., H.W.; Methodology, X.S., W.H., H.X.; Software, X.S., L.C.; Investigation, X.S., W.H., H.X., L.C., H.Z., A.G., N.M. W.W.; Writing – original draft, X.S.; Writing – review & editing, X.S., W.H., H.X., L.C., H.Z., A.G., N.M., H.W. W.W., L.F.; Project Administration, H.W.; Resources, H.W., L.F; Funding acquisition, H.W.

## Additional Information

Supplementary Information is available for this paper.

