## Supplementary Information for "Ultrasound-activated drug release with extracellular vesicles"

**This PDF file includes:**

Figures S1 to S2

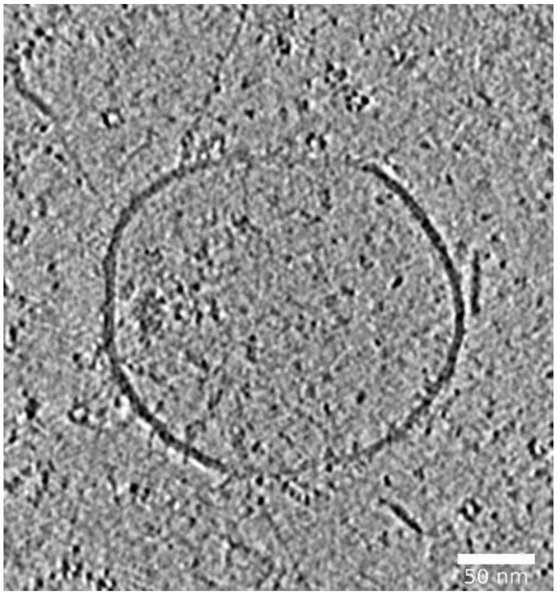


Figure S1. Cryo-Tomography of EVs. Scale bar, 50 nm.

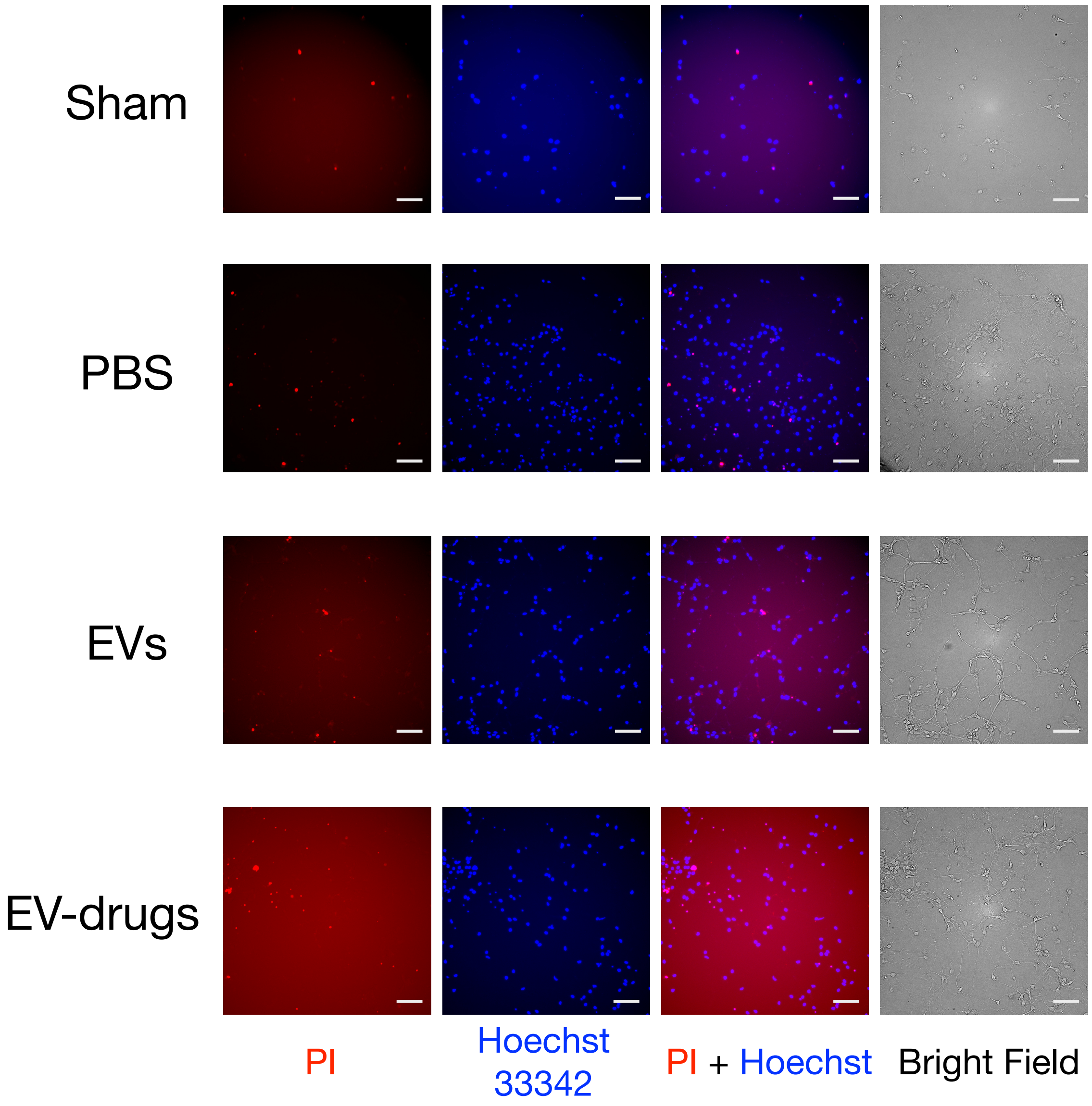


Figure S2. Representative images of toxicity after EV-drugs or the controls groups added to primary cortical neurons for 24 hours.  Propidium Iodide (PI) staining of the dead cells shows red, and the Hoechst 33342 live nucleic acid staining in blue. The scare bar is 100 μm.
